# RIG-I Stimulation Enhances Effector Function and Proliferation of Primary Human CD8 T Cells

**DOI:** 10.1101/2025.01.27.635030

**Authors:** Adham Abuelola Mohamed, Christina Wallerath, Charlotte Hunkler, Gunther Hartmann, Sanda Stankovic, Andrew G Brooks, Martin Schlee

**Author notes:** These authors share senior authorship (Sanda Stankovic, Andrew G Brooks, Martin Schlee).

## Abstract

Cytotoxic CD8 T lymphocytes (CTL) are crucial in antiviral immune responses. However, their recruitment to infection sites renders them at risk of viral infection that could modulate their effector activity. CTL express RIG-I that detects cytosolic viral RNA and subsequently induce antiviral gene expression. Here, we investigated how influenza A virus (IAV) infection influence TCR-dependent effector responses. IAV infection of CTL stimulated RIG-I, and activated downstream pathways including TBK1 and NF-⍰B, resulting in type-I interferon secretion. Transfection of CTL with a pure RIG-I ligand, tri-phosphorylated double stranded RNA(3p-dsRNA), not only stimulated these pathways but also enhanced CTL proliferation *in vitro* and protected them from IAV infection. Analogous with a positive effect on CD8 effector function, both IAV infection and RIG-I ligand transfection enhanced CTL degranulation and cytokine secretion. Conversely, activation of CTL via CD3/CD28-crosslinking increased their susceptibility to IAV infection. Altogether, RIG-I stimulation either by IAV infection or 3p-dsRNA transfection promoted cell intrinsic antiviral pathways and enhanced CD8 effector functions. These findings suggest that RIG-I agonists could enhance and prolong CTL effector function in immunotherapy.

## Introduction

Cytotoxic CD8 T cells are an integral immune component geared to effectively target pathogen-infected or transformed cells (Sun *et al.*, 2023). Their main mode of action is production of pro-inflammatory cytokines such as TNF and IFN-γ which can amplify the immune response (1-3) and cytotoxicity which can directly kill infected cells (4, 5). Due to the nature of TCR-dependent activation which requires direct cell contact (6) between CD8 T cells and infected target cells, there is potential for significant exposure to infectious virions which may in turn impact their capacity to subsequent responses.

CD8 T cells have been shown to be susceptible to infection with a number of viruses. For example, in vitro HHV-6 infection of Jurkat T cells and peripheral blood T cells induced the expression of CD4 protein on mature CD8 T cells, rendering them vulnerable to HIV-1 infection (7, 8). Epstein-Barr virus (EBV), while having a tropism for B cells, has also been found to infect other cells including CD8+ T cells which is clinically obvious in patients with EBV-positive T-cell lymphoproliferative disease(9). Similarly, certain strains of EBV-2 readily infect purified CD8 T cells and induce activation, proliferation, and altered cytokine expression (10). Although HTLV-I primarily infects CD4 T cells, it has been suggested that CD8 T cells may serve as an additional reservoir for HTLV-I in infected patients (11). In addition to DNA virus, negative sense single strand RNA viruses such as measles virus (MeV) and Influenza A virus (IAV) have also been reported to infect CD8 T cells. Peripheral blood mononuclear cells (PBMC) of prodromal measles patients were shown to be infected by MeV with specific tropism for CD8+ T cells (12) and *in-vitro* experiments demonstrated that MeV can replicate in T lymphocytes, including CD8 T cells (13). In mouse models, lymphocytic choriomeningitis virus (LCMV) can productively infect CD8 T cells and influenza A virus (IAV) has also been observed to directly infect CD8 T cells (14). Consequently, there is potential for T cell intrinsic anti-viral mechanisms to significantly impact not only the susceptibility of T cells to viral infection but also their subsequent TCR-dependent responses.

Autonomous antiviral defence mechanisms have been described in immune cells with substantial impact on their effector response. Activation of these innate nucleic acid receptors typically triggers a robust antiviral immune response, characterized by the secretion of type I IFN (IFN-I), cytokines, and chemokines. Toll-like receptors (TLRs) 3, 7, 8, and 9 sense viral nucleic acids within endosomes whereas cytosolic RNA and DNA is sensed by retinoic acid-inducible gene I (RIG-I)-like helicases (RLH) and cyclic GMP-AMP synthase (cGAS) respectively (15-18). There is a wealth of data showing that ligand recognition by TLRs expressed on APC plays a crucial role in T cell activation (19) but the role of T cell intrinsic recognition of nucelic acids is less well described. Direct stimulation of TLR3 expressed on CD8 T cells resulted in increased IFN-γ production without affecting their cytotolytic function or proliferative capacity (20). Similarly, *in vitro* analyses demonstrated that direct exposure of human CD8 T cells to TLR7 ligands directly increased the expression of activation markers CD69 and CD25, along with the production of IFN-γ in (21).

Few studies have directly addressed the potential of ligand recognition by T cell intrinsic RNA helicases to modulate T cell function. RIG-I is probably the best described of these helicases and specifically recognizes structural RNA motifs that are a common feature of viral RNA or replicative intermediates that are generated during viral infection(16, 22-24). RIG-I signals through the activation of the mitochondrial antiviral signaling protein (MAVS), which activates IkappaB kinases (TBK1/IKKα/IKKβ/IKKε) which in turn activate transcription factors such as nuclear factor kappa B (NF-⍰B ) and IRF3/7 (17). In macrophages and dendritic cells, this typically results in the secretion of type-I and type-III interferons and cytokines such as IL-27 (25). The secreted IFN-I then signals through the IFN-α/β receptor complex (IFNAR) to activate Janus kinase 1 (Jak1) and Tyrosine kinase 2 (Tyk2), which subsequently phosphorylate signal transducer and activator of transcription 1 (STAT1) and STAT2 proteins (26). Phosphorylated STAT2 in association with STAT1 and interferon regulatory factor 9 (IRF-9) forms the interferon-stimulated gene factor 3 (ISGF3) (24). Once translocated to the nucleus, it activates the transcription of a range of antiviral effector genes, including the interferon-induced protein with tetratricopeptide repeats 1 (IFIT1) (17, 27). Synthetic ligands such as short (>/=20bp) 3p-dsRNA mimic viral RNA and are potent and specific inducers of the RIG-I signaling pathway (16).

While RIG-I deficiency has been shown to diminsh T cell responses following IAV infection where deficient mice exhibited lower level of activated CD8 T cells (28) it is unclear whether this is due to direct effects on RIG-I expressed by CD8 T cells themselves or rather reflects indirect effects such as reduced activation or cytokine production by APCs. This study investigated the impact of CD8 T cell intrinsic activation of RIG-I on the effector function by either direct infection of CD8 T cells with IAV or via the transfection of pure RIG-I ligands (3p-dsRNA) into the cytosol of CD8 T cells. The data show that in vitro IAV infection induced RIG-I and downstream TBK1 and NF-⍰B pathways, as well as the secretion of type-I IFN in infected cells. Secondly, although activated CD8 T cells were more succeptible to IAV infection, their effector functions such as degranulation and the secretion of IFN-γ and TNF, were enhanced by IAV infection. Moreover, targeted activation of RIG-I with 3p-dsRNA drove similar responses, namely increased release of type-I IFN and enhanced degranulatioin and IFN-γ secretion by CD8 T cells. Importantly, this RIG-I activation provided protection against subsequent IAV infection and enhanced the proliferative capacity of the cells. Overall, these findings demonstrate that RIG-I activation in CD8 T cells has the potential to enhance the effector function of these cells and offers a new avenue to improve the outcomes of T cell therapies.

## Results

### 1. Activated CD8 T cells are more susceptible to IAV infection

To assess the impact of exposure to IAV on CD8 T cell function, resting or *in vitro* activated CD8 T cells were initially infected with IAV for 8h, after which the proportion of NP+ cells was determined using flow cytometry. The proportion of NP+ cells was significantly higher following anti-CD3/CD28-stimulation (16% vs 7%), (Figure 1A & B). The differences in these proportion of NP+ cells were not attributable to differences in cell viability as this was not significantly different between the groups at 25h post-infection (Supp. Fig. 1A). Moreover, although there was clear evidence of infection at 25h post-infection, the absence of newly released infectious virus in the supernatant indicated that the infection was non-productive (Supp. Fig. 1B).

**Figure 1.**
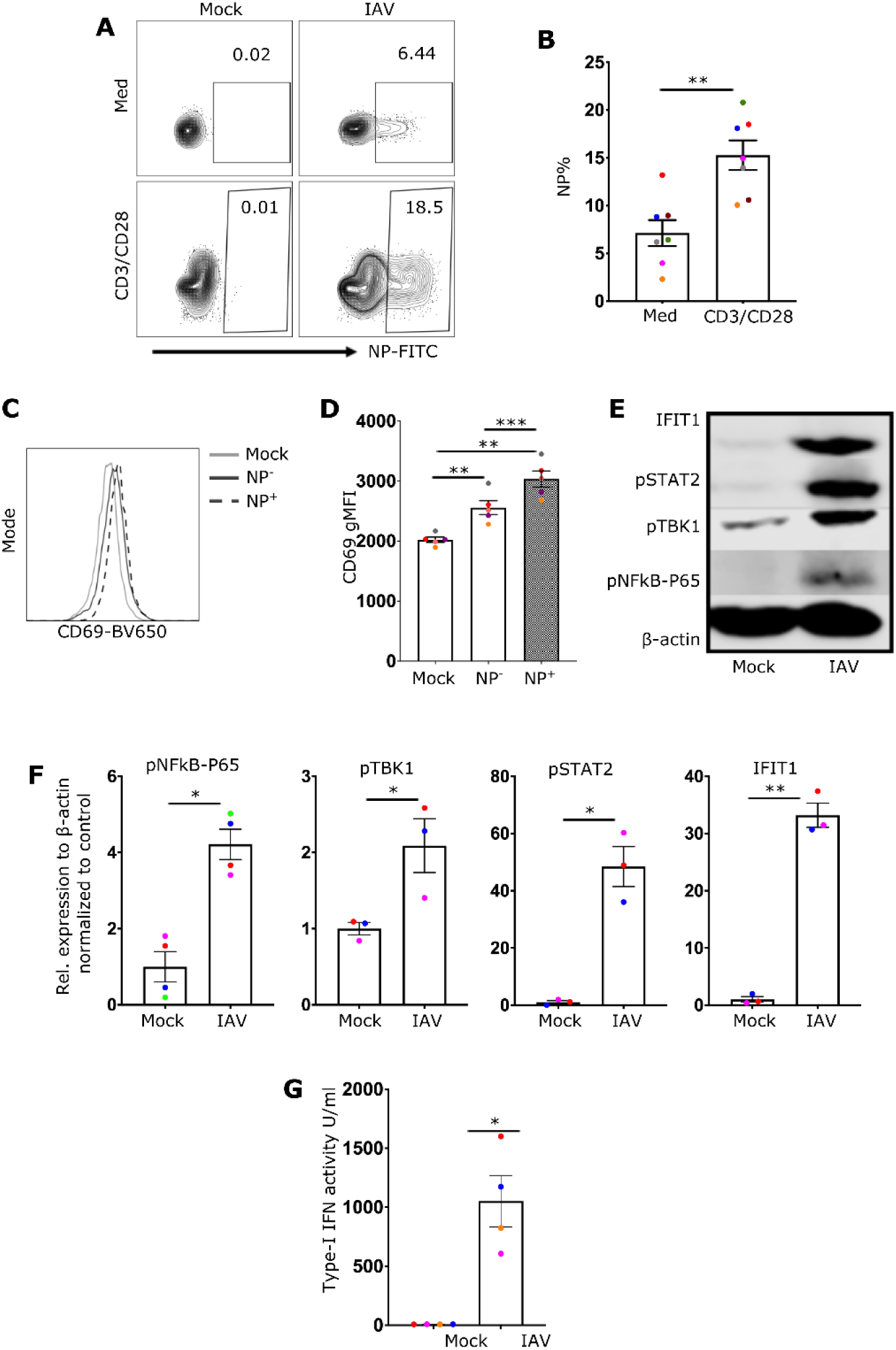
Infection of human primary CD8 T cells leads to the activation of NFkB, TBK1 and interferon response. **(A)** Representative flow cytometry plots for CD8 T cells cultured for 3 days with or without anti-CD3/CD28 antibodies, then treated with either media (mock) or infected with 10 MOI of IAV for 8hrs. **(B)** a bar chart summarizing the results for 7 different donors infected with IAV with or without anti-CD3/CD28 antibodies. **(C)** representative histogram of CD69 expression in mock-treated, NP-, and NP+ cells from the infection condition. **(D)** Quantification of CD69 gMFI. **(E)** Western blot image of mock- and IAV-infected CD8 T cells, illustrating different proteins associated with nucleic acid receptor stimulation (pNFkB-P65 and pTBK1) and interferon response (IFIT1 and pSTAT2). **(F)** Quantification of the proteins illustrated in E. **(G)** a bar chart demonstrating the IFN-I activity detected by TBK1^-/-^ IKKα^-/-^ IKKβ^-/-^ & IKKε^-/-^ THP1 dual reporter cells. Every donor is represented by a colored dot, bars show mean ± SEM. Paired t-test was used for two group comparison and one-way ANOVA followed by Dunnett’s correction for more than two groups (*p < 0.05, **p < 0.01, and ***p < 0.001)

### 2. IAV drives RIG-I, NF-⍰B, TBK1 activation and type-I IFN response in CD8 cells

The next objective was to assess if CD8 T cells undergo activation in response to IAV by measuring CD69 expression, an early indication of cellular activation. IAV exposed CD8 T cells exhibited slightly increased levels of CD69 expression, relative to mock treated cells. Moreover, amongst IAV-exposed cells, NP+ cells had higher levels of CD69 expression than the NP-population (Figure 1C, D).

As IAV infection can activate nucleic acid sensors, potentially driving the production of cytokines, the impact of IAV infection on different pathways downstream of nucleic acid receptors was assessed. Protein immunoblotting revealed that IAV infection induced phosphorylation of both NF-⍰B.p65 and TBK1 (Figure 1E, F). Additionally type-I IFN was detected in the supernatants of infected CD8 T cells 24 hours post-infection (Figure 1G). Type-I IFN can drive an autocrine signaling cascade by binding to IFNARs which results in STAT2 phosphorylation. Consistent with signaling through IFNARs, immunoblotting showed STAT2 phosphorylation in IAV-infected-but not mock treated CD8+ T cells (Figure 1E, F). In line with the production and recognition of IFN-I, the expression of IFIT1, an interferon-induced protein was also strongly induced in IAV-exposed cells (Figure 1E, F). Although phosphorylation of NF-⍰B may be driven by exposure to exogenous cytokines, the activation of TBK1 and the release of type-I IFN in IAV-infected CD8+ T cells demonstrates the involvement of upstream nucleic RNA receptors, such as RIG-I in the activation process.

To better identify the role of RIG-I receptors in the CD8 T cell IFN response during viral infection, CRISPR-editing was used to delete RIG-I or STAT2 from primary CD8 T cells. As a key downstream component of IFNAR, STAT2 is crucial for the autocrine amplification loop that enhances T cell activation. Investigating this could clarify the role of the amplification loop. The efficiency of the knockout was evaluated by analyzing protein expression relative to beta-actin. Immunoblot analyses showed the expression of both RIG-I (Figure 2A,B) and STAT2 (Figure 2A,D) proteins was reduced by more than 50% in the CRISPR-edited cells. Moreover, the expression of IFIT1 following IAV was essentially abrogated in both RIG-I and STAT2 knock out cells, demonstrating the importance of both RIG-I and STAT2 signaling in the interferon response through autocrine activation of IFNAR (Figure 2C, E).

**Figure 2.**
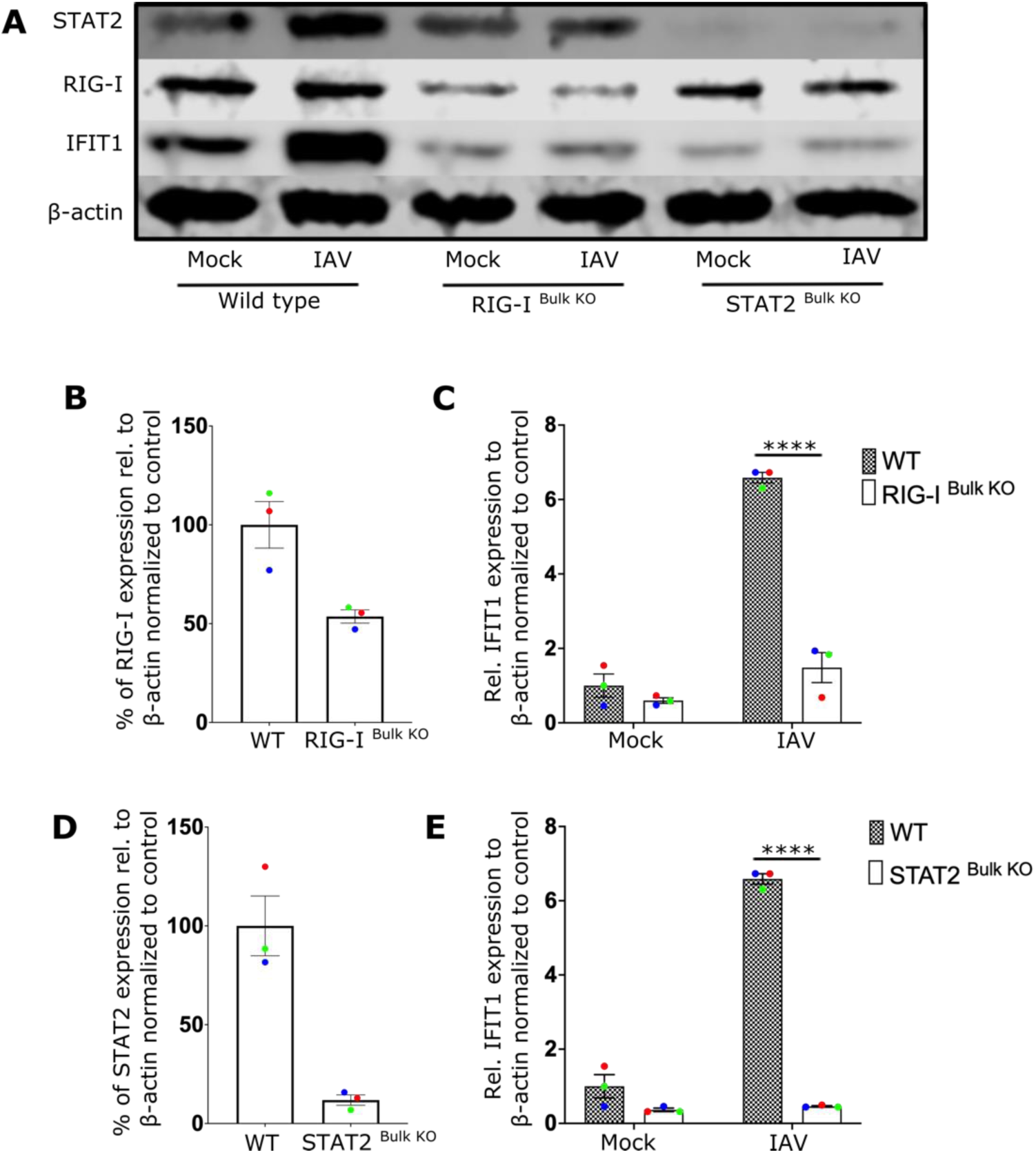
IAV infection triggers antiviral IFN response in CD8 T cells via RIG-I and STAT2 signaling pathways. **(A)** Representative Western blot image of wildtype (WT), RIG-I ^Bulk^ ^KO^, and STAT2 ^Bulk^ ^KO^ CD8 T cells treated with media (Mock) or infected with influenza A virus (IAV). **(B, D)** show the relative expression level of RIG-I and STAT2 respectively in WT and knockout cells. **(C, E)** displays the relative expression level of IFIT1 in WT cells and RIG-I ^Bulk^ ^KO^ or STAT2 ^Bulk^ ^KO^ cells treated as mentioned. Every donor is represented by a colored dot, bars show mean ± SEM. Two-way ANOVA followed by Bonferroni’s correction for more multiple comparison (****p < 0.0001).

### 3. IAV infection increases the effector function of CD8 T cells

The observation that IAV infection of CD8 T cells activated nucleic acid sensing pathways resulting in both activation of NF-⍰B and the secretion of IFN-I, suggested that it also had the potential to modulate effector functions. To assess this more formally, CD8 T cells were cultured in plates coated with or without anti-CD3 and anti-CD28 for three days then infected with IAV for 5 hr. After infection, the degranulation was assessed by staining for CD107a and cytokine production by staining for intracellular IFN-γ. As expected in the absence of stimulating mAb, neither mock-treated nor IAV-infected CD8 T cells exhibited notable degranulation or IFN-γ production (Figure 3 A-C). Following stimulation, both the degranulation and IFN-γ responses of IAV-exposed but NP-CD8 T cells were similar to those of mock treated controls (Figure 3D-F). However, the responses of NP+ CD8 T cells were significantly elevated (Figure 3E,F), suggesting an intrinsic impact of IAV in enhancing CD8 T effector function, beyond simply exposure to secreted IFN-I (Figure 3).

**Figure 3.**
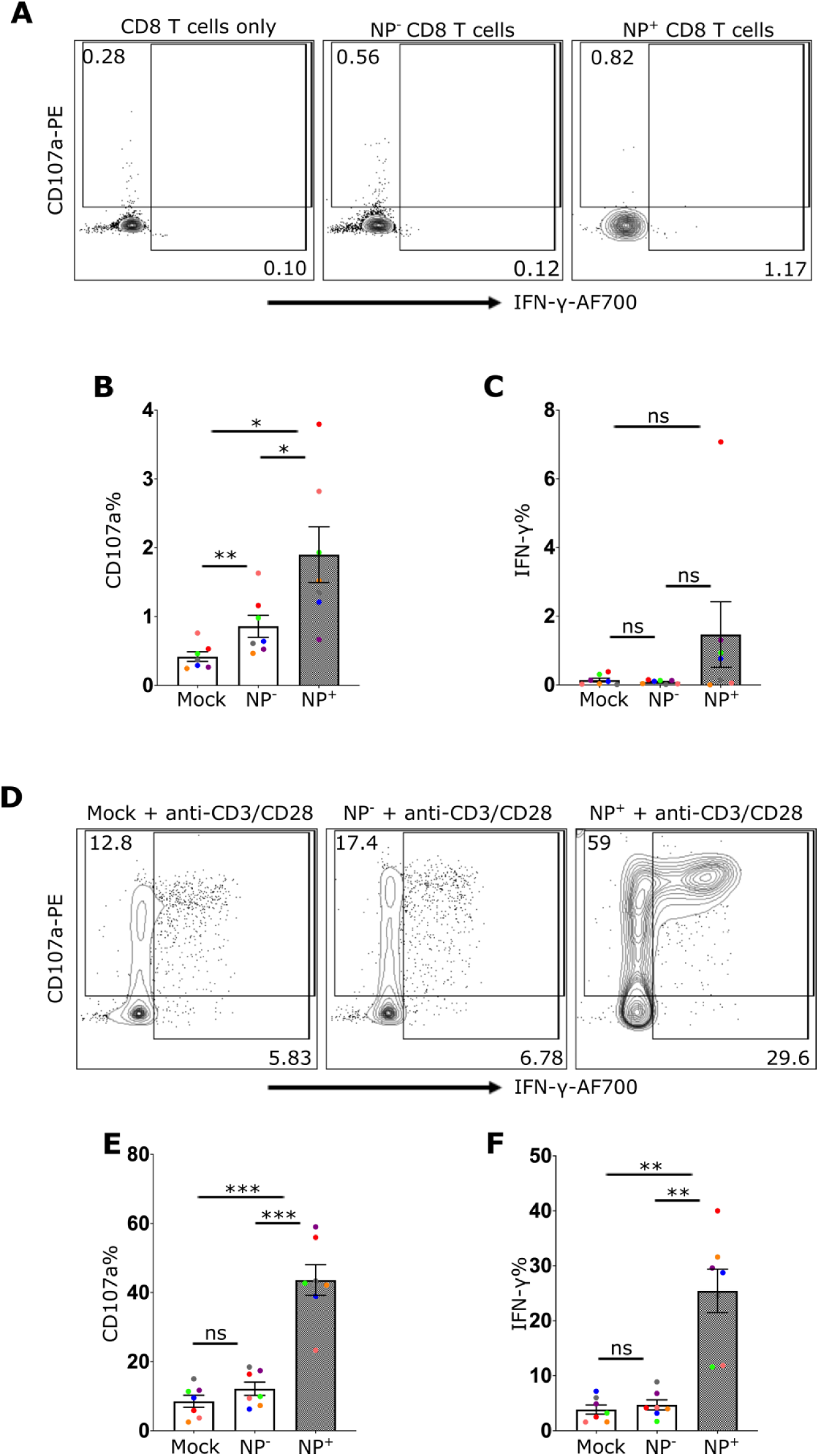
Enhanced Effector Function in CD8 T Cells upon IAV Infection. Flow cytometry analysis of CD8 T cells 5 hrs post-infection with or without activation. **A** and **D** display the representative dot plots of CD107a and IFN-γ expression in CD8 T cells without anti-CD3/CD28 antibody activation **(A)** or with anti-CD3/CD28 antibody activation **(D)**. Cells were treated with media control (Mock) or infected with IAV, gated based on the expression of viral nuclear protein: IAV-exposed cells (NP-) or infected cells (NP+). **(B)** and **(C)** show the frequencies of CD107a+ **(B)** or IFN-γ+ **(C)** CD8 T cells without activation, while **(E)** and **(F)** show the frequencies with antibody activation. Every donor is represented by a colored dot, bars show mean ± SEM. Repeated measure one-way ANOVA followed by Dunnett’s correction for more than two groups (ns= not significant, *p < 0.05, **p < 0.01, and ***p < 0.001).

### 4. 3p-dsRNA triggers the activation of TBK1 and NF-⍰B pathways in CD8 T cells resulting in the induction of type-I interferon

While the data clearly showed that IAV-infection could activate RIG-I activation to drive IFN-I production, the enhanced effector responses could potentially be attributed to broader effects of IAV infection on CD8 T cell function. Consequently, to better target RIG-I a specific in vitro generated RNA ligand, 3p-dsRNA, or control RNA (that does not activate RIG-I) was introduced into CD8 T cells. Flow cytometry using FAM-labelled 3p-dsRNA confirmed the successful delivery of the RIG-I ligand into CD8 T cells, as indicated by FAM+ cells (Suppl. Fig. 2A). Similar to the response observed with IAV, the introduction of 3p-dsRNA induced the upregulation of the activation marker CD69 (Figure 4A, B), phosphorylation of NF-κB-p65 and TBK1 (Figure 4C, D), type-I IFN secretion (Figure 4E), and phosphorylation of STAT2 downstream of IFN-I receptors (Figure 4D), ultimately leading to the induction of the interferon-stimulated protein IFIT1 (Figure 4C, D).

**Figure 4.**
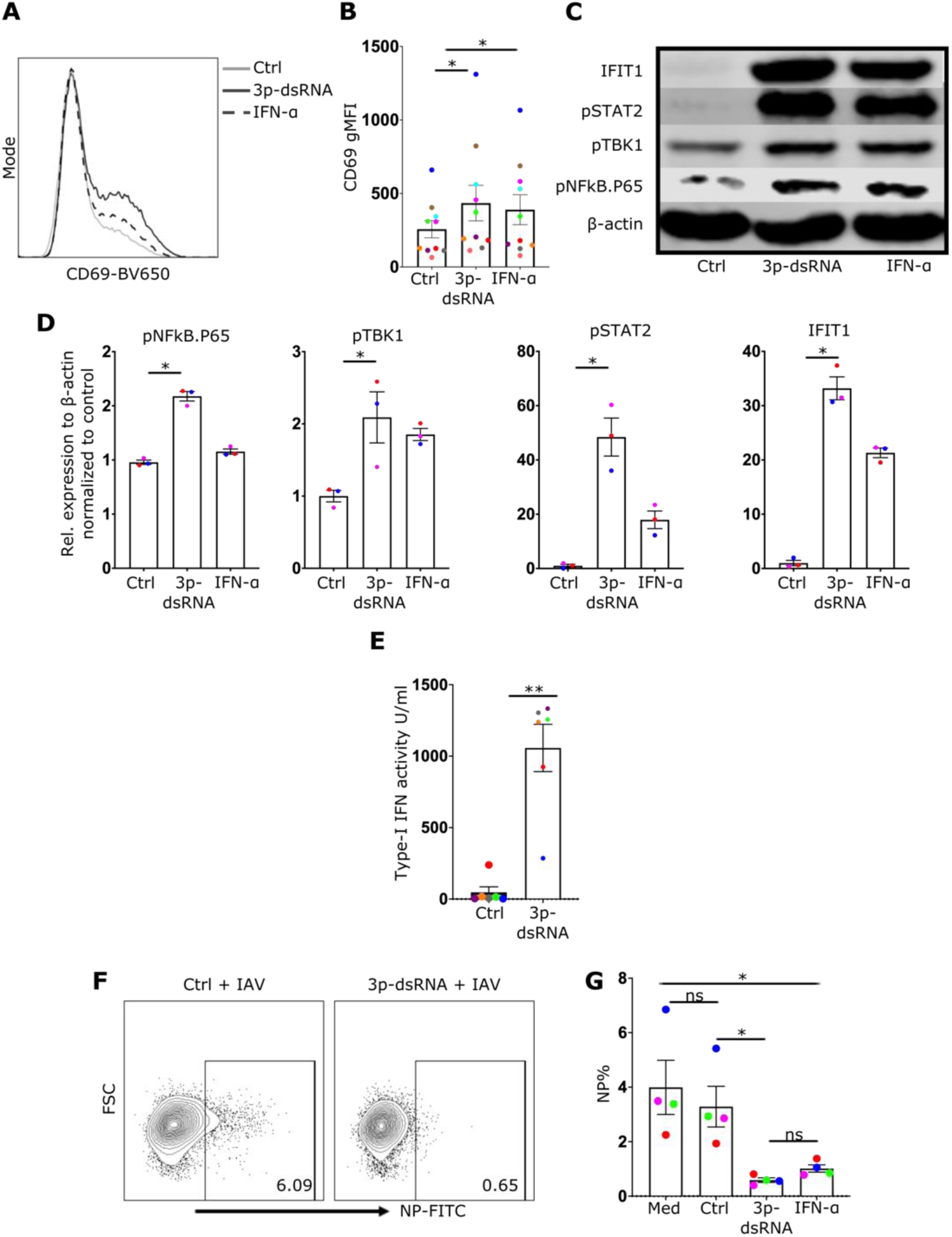
RIG-I stimulation in CD8 T cells stimulates same pathways as in IAV infection and inhibits subsequent IAV infection. **(A)** An illustrative histogram displaying CD69 expression in CD8 T cells that were treated with control (Ctrl), RIG-I ligands (3p-dsRNA) or IFN-α. **(B)** Quantification of CD69 gMFI for 10 different donors. **(C)** Western blot image showing proteins from CD8 T cells that were treated with the same conditions as mentioned in **A**. **(D)** Bar charts illustrating the relative levels of listed proteins to β-actin in CD8 T cells (n=3). **(E)** Bar chart shows the measurement of IFN-I activity using THP1 reporter cells. **(F)** Flow cytometry plots showing CD8 T cells treated as mentioned and then infected with IAV for 8 hrs. **(G)** Bar chart showing the frequencies of infected cells (NP+) for CD8 T cells treated as labelled. Every donor is represented by a colored dot, bars show mean ± SEM. Paired t-test was used for two group comparison and repeated measures one-way ANOVA followed by Dunnett’s correction for more than two groups (ns= not significant, *p < 0.05, **p < 0.01).

Not only did RIG-I stimulation enhance activation of CD8 T cells, pre-treatment with RIG-I ligand (or IFN-I) in contrast to negative control RNA, protected NP+ CD8 T cells from IAV infection (Figure 4F, G).

As observed following IAV-infection, following transfection of 3p-dsRNA, the induction of IFIT1 was reduced by 5-fold in RIG-I- and by 6-fold in STAT2-deficient CD8 T cells relative to unmodified T cells (Figure 5). further confirming that RIG-I stimulation is upstream of NF-⍰B and TBK1 pathways that led to type-I IFN secretion in CD8 T cells. Furthermore, the addition of neutralizing antibodies to block IFNAR2 resulted in the reduction of STAT2 phosphorylation and IFIT1 protein expression, confirming that both effects depend on the autocrine/paracrine effects of secreted IFN-I (Supp. Fig. 2B).

**Figure 5.**
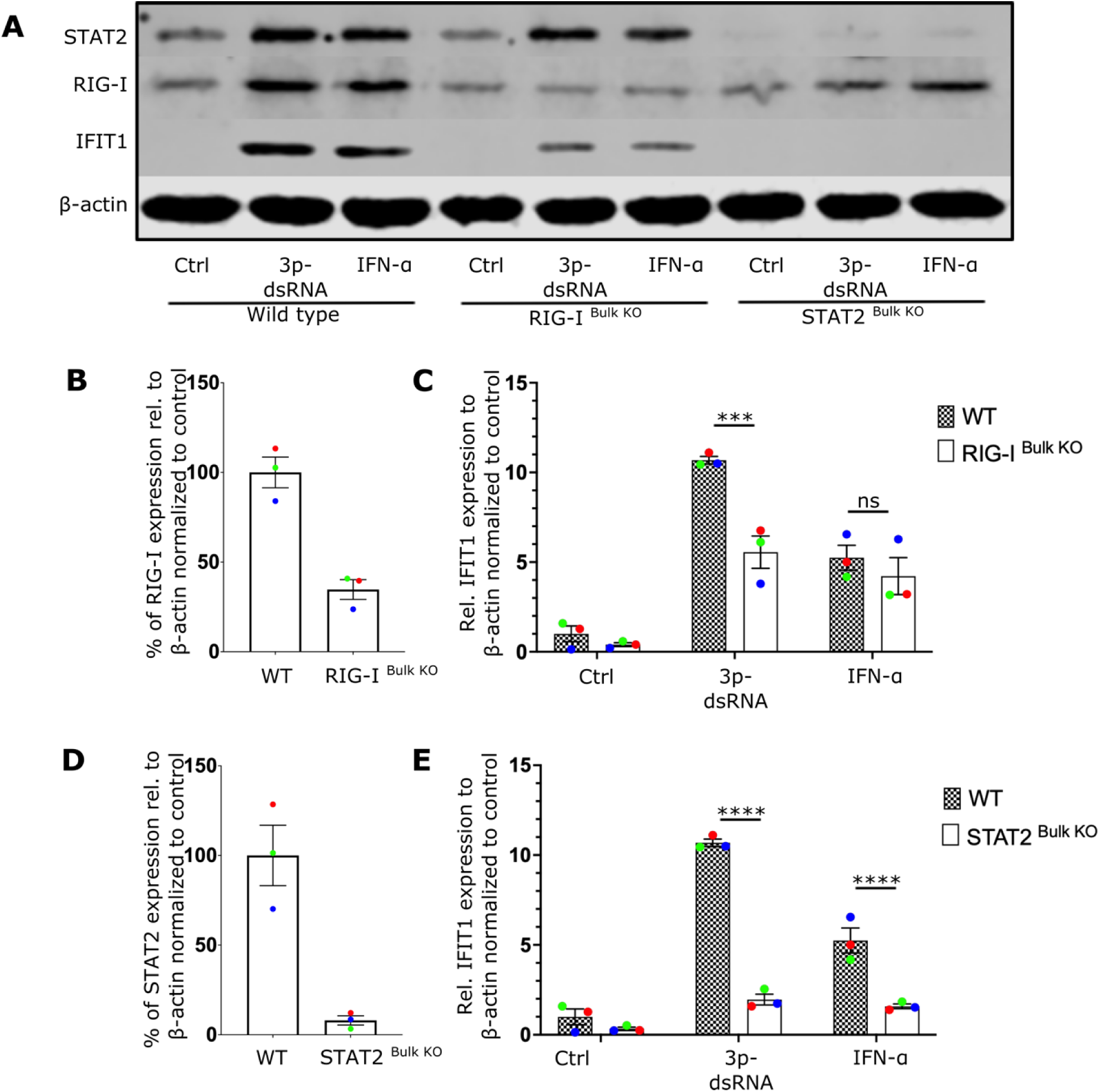
RIG-I ligands trigger interferon response in CD8 T cells via RIG-I and STAT2 pathways. **(A)** Representative Western blot displays the expression of IFIT1, RIG-I, and STAT2 proteins in CRISPR/Cas9 genetically edited CD8 T cells treated as described. The cells included wildtype (WT), RIG-I ^Bulk^ ^KO^, as well as STAT2 ^Bulk^ ^KO^ cells. **(B, D)** Bar chart showing the relative expression level of RIG-I and STAT2 respectively. **(C, E)** The bar charts display the relative expression level of IFIT1 in RIG-I ^Bulk^ ^KO^ cells and STAT2 ^Bulk^ ^KO^ respectively. Every donor is represented by a colored dot, bars show mean ± SEM. Two-way ANOVA followed by Bonferroni’s correction for multiple comparisons (ns= not significant, ***p < 0.001, and ****p < 0.0001).

### 5. RIG-I ligands boost the function of CD8 T cells

To determine whether the enhanced effector response of infected CD8 T cells was directly attributable to the RIG-I pathway CD8 T cell effector function, CD8 T cells were treated with control RNA, 3p-dsRNA or IFN-α and their response to antibody activation was evaluated. In line with the observations during IAV infection, a significant increase in degranulation (Figure 6A,B), as well as in the production of IFN-γ (Figure 6A,C) and TNF (Figure 6D), was observed after exposure to 3p-dsRNA. On the other hand, IFN-α led to enhanced effector function compared with the control but to a lesser extent than RIG-I ligands. Additionally, enhanced effector function in response to antibody activation was observed in sorted CD8 T cells (Supp. Fig. 3), ruling out the potential effects of contaminating myeloid cells. These findings suggest that the increased activation of CD8 T cells following exposure to 3p-dsRNA was due to intrinsic activation of the RIG-I pathway within CD8 T cells themselves. In summary, it can be concluded that CD8 T effector function is significantly enhanced by IAV and 3p-dsRNA in a RIG-I dependent mechanism.

**Figure 6.**
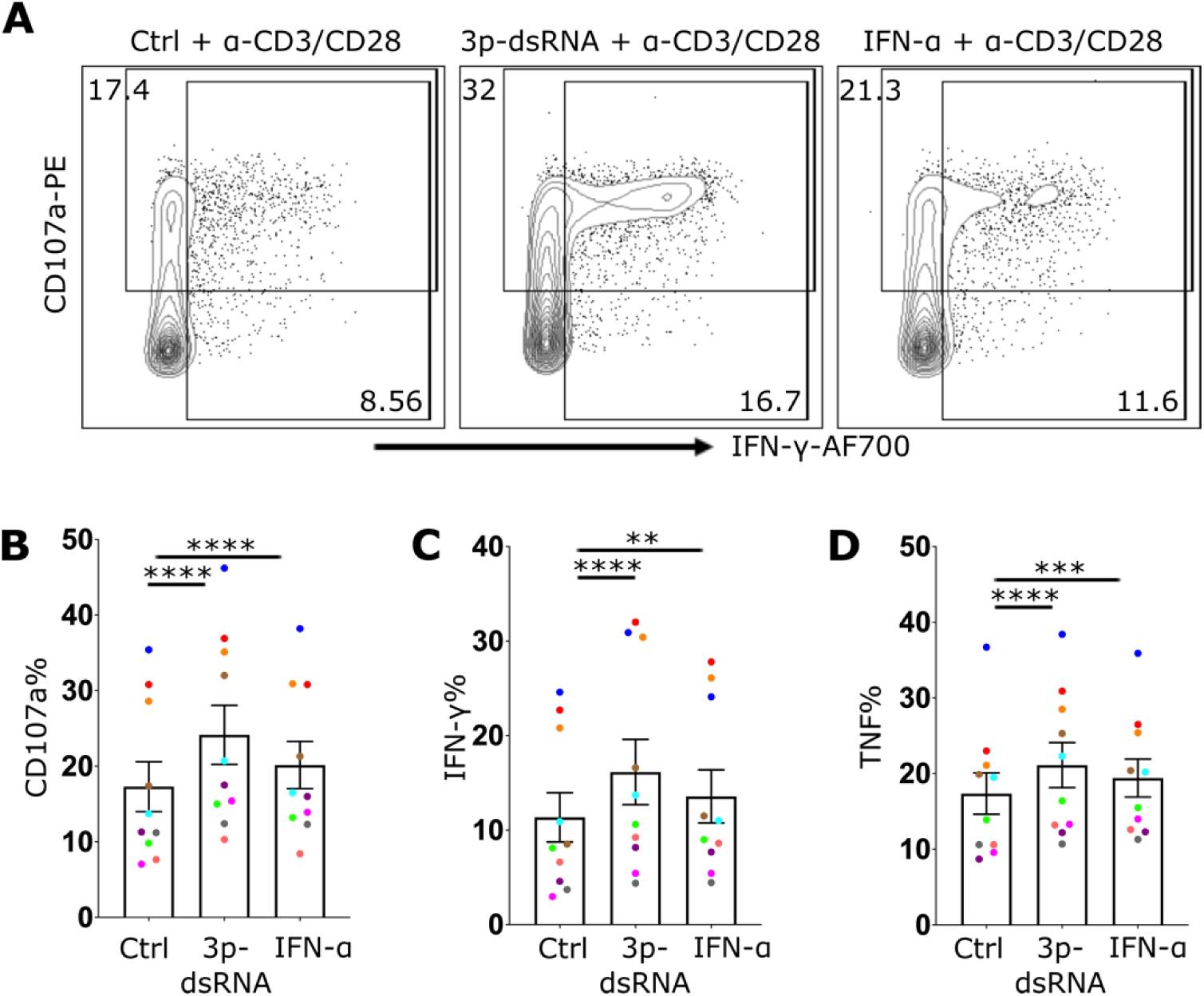
RIG-I stimulation in CD8 T cells enhances their effector function. **(A)** Flow cytometry plot showing CD8 T cells treated with control RNA (Ctrl), 3p-dsRNA or IFN-α overnight followed by anti-CD3/CD28 activation for 4 hrs. The plot displays the expression of CD107a and IFN-γ. **(B-D)** Bar charts illustrating the frequencies of CD107a+ **(B)**, IFN-γ+ **(C)** and TNF+ **(D)** CD8 T cells after antibody-mediated activation. Every donor is represented by a colored dot, bars show mean ± SEM. Repeated measures one-way ANOVA followed by Dunnett’s correction for more than two groups (ns= not significant, **p < 0.01, ***p < 0.001, and ****p < 0.0001).

### 6. RIG-I activation enhances proliferation of CD8 T cells

RIG-I stimulation enhanced TCR-mediated responses which suggested it may influence not only activation and effector response but also proliferation. We compared the proliferation of RIG-I stimulated CD8 T cells with 3p-ssRNA (negative control) and IFN-α, which is known to activate these cells (29). The CD8 T cells stimulated through RIG-I exhibited an enhanced proliferation compared to negative control or IFN-α. This was shown by the higher percentages of proliferating cells (cell trace violet low) (Figure 7A) and the higher proliferation ratio based on cell counts (Figure 7B). While CD25 expression remained similar across all conditions, the expression of the activation marker CD69 was significantly increased in RIG-I- and IFN-α-stimulated CD8 T cells (Figure 7C, D), with RIG-I ligands showing the highest expression levels. This concludes that RIG-I stimulation may have a beneficial effect on the prolonged activation of CD8 T cell leading to enhanced cell expansion.

**Figure 7.**
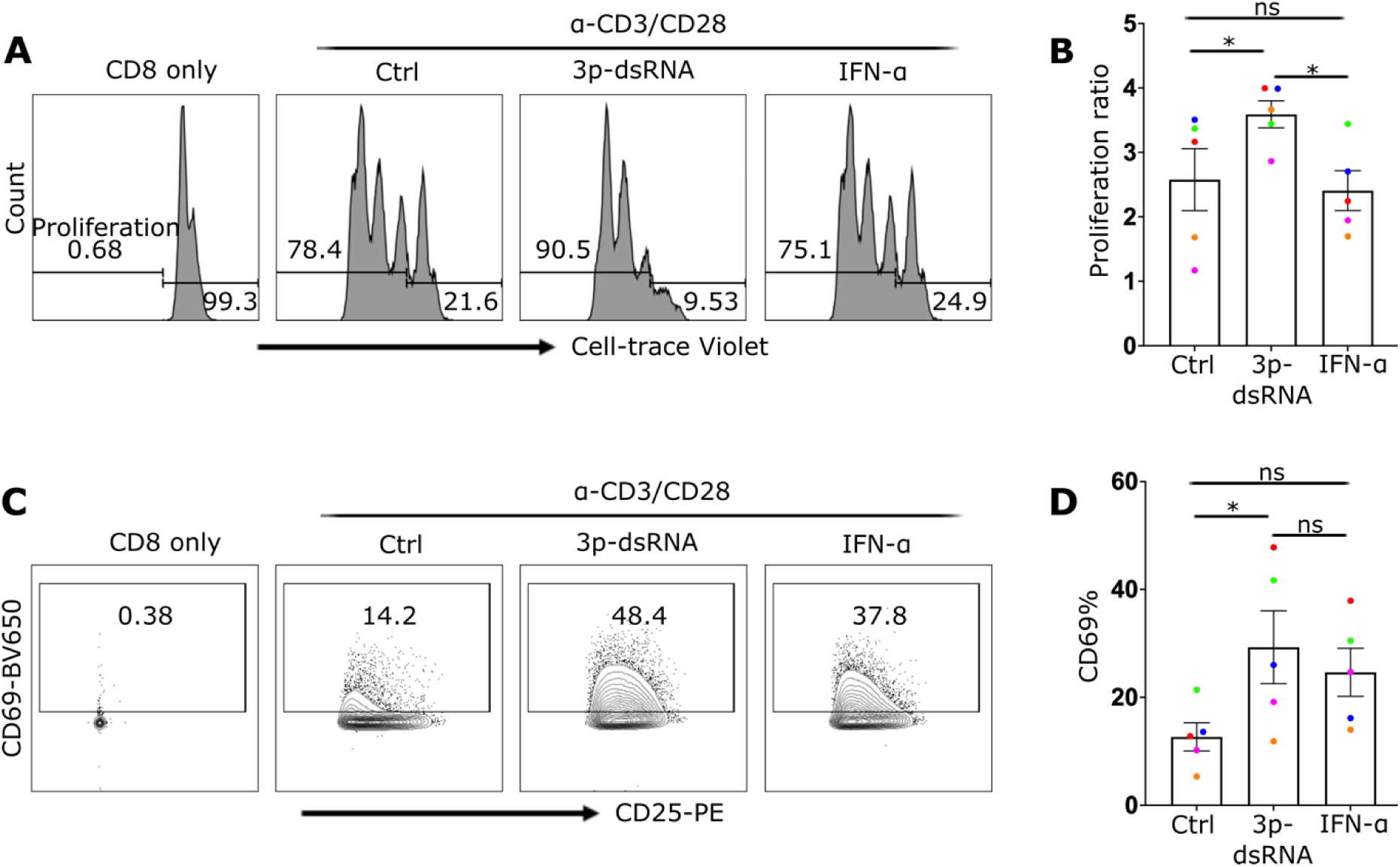
RIG-I ligands enhance CD8 T cell activation and proliferation. **(A)** Histograms showing CD8 T cells with conditions as labelled. The leftmost peak represents the control group with no stimulation or proliferation. The other peaks represent cells stimulated overnight with control RNA (Ctrl), 3p-dsRNA, or IFN-α, followed by a 3-day culture in CD3/CD28 coated plates. The histogram demonstrates the proliferation tracking dye (Cell-trace Violet), with lower dye intensity indicating higher proliferation. **(B)** The proliferation ratio of each condition calculated by dividing the total cell count of the anti-CD3/CD28 treated cells by cells incubated with no anti-CD3/CD28. **(C)** Flow cytometry plots for the same conditions as in **A**, displaying CD69 on the Y axis and CD25 on the X axis. **(D)** presents the CD69 frequencies. Every donor is represented by a colored dot, bars show mean ± SEM. Repeated measures one-way ANOVA followed by Dunnett’s correction for more than two groups (ns= not significant, *p < 0.05).

## Discussion

During the process of targeting infected cells, immune cells such as CD8 T cells often encounter infectious virions, especially at synaptic junctions, potentially increasing their vulnerability to infection (30). The direct effect that this process can have on CD8 T cell effector function has not been thoroughly explored. Previous studies have highlighted the significant indirect impact of ligand recognition by nucleic acid receptors on CD8 T cell responses (28, 31, 32) through enhanced activation of antigen-presenting cells such as dendritic cells, leading to the enhanced secretion of immunomodulator cytokines and expression of co-stimulating molecules (33). There is however evidence that T cell intrinsic activation of nucleic acid receptors can significantly alter their function. For instance, TLR3 receptor stimulation in murine CD8 T cells resulted in increased IFN-γ production, without affecting their cytolytic or proliferative activity (20). Similarly, *in vitro* studies have shown that TLR7 ligands can directly amplify the activation and cytokine production of human CD8 T cells (21). By contrast, stimulation of the DNA receptor cGAS/STING pathway induced IFN-I but negatively affected T cell functions by increased cell death and decreased proliferation (34).

RIG-I has been proposed to have therapeutic potential by enhancing the effector function of CD8 T cells. Administration of RIG-I ligands into mice synergistically complemented the outcome of checkpoint inhibition, leading to the activation and expansion of antigen-specific CD8 T cells *ex vivo* and enhancing their antitumor response *in vivo (35)*. However, the role of intrinsic activation of RIG-I in CD8 T cells was not addressed.

On the other hand, although CD8 T cells are not the primary viral targets, evidence in both human and mouse systems suggests their susceptibility to viral infections, including IAV (7, 8, 12, 14, 36-38). Our first observation that activated CD8 T cells are more susceptible to IAV infection, makes this notion even more relevant clinically. In line with our study, previous studies have shown that both IAV and HTLV-I viruses tend to infect activated human lymphocytes preferentially over resting cells (39, 40). This increased susceptibility might be attributed to distinct glycosylation profiles, with activated cells expressing higher levels of sialyl glycans that potentially facilitate greater IAV entry (41). Moreover, the increased metabolic activity in activated cells could potentially promote viral infection and replication, leading to increased viral protein expression (42). Conversely, resting cells may have a more robust intrinsic antiviral response, which could explain the lower susceptibility of resting CD8 T cells to infection. This is analogous to research showing that resting CD4 T cells are less susceptible to HIV infection compared to their activated counterparts (43). Alternatively, upon infection, naïve cells might experience rapid cell death, resulting in a lower proportion of infected cells in our flow cytometry data. Further investigation is necessary to better understand the underlying reasons for the increased proportions of NP+ cells observed upon activation of CD8 T cells.

Our results indicate that the most distinct effect of IAV infection is via activation of RIG-I pathway. This not only enhances CD69 expression, but activates downstream TBK1 and NF-κB pathways, and IFN-I secretion. Furthermore, we used primary CRISPR/Cas9-edited CD8 T cells to analyse the signalling pathways. The induction of the IFN-I response in CD8 T cells following either IAV infection or 3p-dsRNA transfection was dependent on RIG-I. Although IFIT1 is known to be directly regulated by RIG-I receptor signalling through IRF3 (44), we observed that deleting STAT2 downstream of the IFNAR nearly abolished the induction of IFIT1 expression in CD8 T cells, indicating a crucial role of the autocrine IFN feedback/amplification loop via the IFNAR in CD8 T cells for the full induction of the antiviral response.

Functionally, RIG-I activation had a co-stimulatory effect on TCR-mediated activation. This was mainly observed for infected cells (NP+) but not for non-infected bystander cells, demonstrating strong dependence on the intracellular RIG-I signalling pathway, rather than being dependent on secreted IFN-I from these cultures. Additionally, treatment with IFN-α had a lesser impact on CD8 T cell effector function compared to the effects of RIG-I ligands, further supporting that the synergy between RIG-I and TCR-mediated signalling depends on more RIG-I induced signalling molecules and genes than solely IFN-I. These results highlight the potential of RIG-I receptor signalling to amplify CD8 T cell function, suggesting its potential as a therapeutic target for enhancing therapies based on CD8 T cells. An important consideration is that RIG-I signalling not only protects CD8 T cells from infection and enhances their effector function, but our results also indicate a strong positive effect on the proliferation of CD8 T cells. This was unexpected, since the stimulation of the cGAS/STING pathway that shares parts of the cytosolic signalling with the RIG-I pathway inhibits T cell growth and therefore impedes T cell effector function (34).

NF-κB signalling may be the key pathway responsible for the activating effects of RIG-I as NF-κB it is activated by both RIG-I (45) and TCR-(46) signalling but not by IFNAR.

This study sheds light on the impact of inherent RIG-I pathway activation on cellular responses during viral infections, particularly with IAV. The findings demonstrate that RIG-I activation can enhance TCR-dependent activation of CD8 T cells, affecting processes such as degranulation, proliferation, cytokine release, and providing defense against subsequent IAV infections. Additionally, we show that CD8 T cells are capable of secreting IFN-I, induced by direct RIG-I ligand transfection or by IAV infection. This reveals an expanded role for these cells in combating viral infections and immune challenges. Beyond their direct antiviral and anticancer effects, CD8 T cells’ ability to secrete IFN-I enables them to stimulate other immune cells and alert non-infected cells to the viral threat, thereby enhancing the overall immune response.

Taken together, these results suggest the potential therapeutic use of RIG-I activation to enhance protective immune responses against viral pathogens. More extensive research is needed to comprehensively investigate the therapeutic implications and real-life effects of RIG-I activation on antiviral immunity. A deeper understanding of how viral infections interact with the adaptive immune response, in particular CD8 T cell-mediated immunity, could lead to innovative immunotherapeutic approaches. These approaches aim to effectively combat viral or other microbial infections and enhance the immune response against tumours. One of the applications of T cell therapy is CAR T cell therapy. CAR T cells have revolutionized the treatment of certain cancers but still faces limitations, including lengthy manufacturing times, suboptimal persistence during and after ex vivo cultivation, and reduced efficacy against solid tumors (47). RIG-I stimulation, which enhances T cell proliferation, offers a promising approach to overcome these challenges. By boosting T cell expansion, RIG-I stimulation could potentially shorten the manufacturing process, improve the persistence of CAR T cells during cultivation and after reinfusion, and enhance their effectiveness against solid tumors. This could make CAR T cell therapy more accessible and effective, addressing some of the current hurdles in its application.

Since the last decade, the cytosolic dsDNA sensor cGAS and the signaling hub STING downstream of cGAS on one hand and RIG-I like helicases on the other hand have been targeted in anti-tumor therapy, reviewed in (48, 49). Our data suggest that RIG-I ligands have an important advantage over cGAS/STING agonists as they are able to directly enhance CD8 T cell effector function and proliferation.

## Materials and Methods

Mononuclear cells from the peripheral blood (PBMCs) of healthy individuals were collected through a density gradient centrifugation process after receiving written consent and approval from the relevant institutional review board. CD8 T cells were purified from the Isolated PBMCs using the EasySep™ Human CD8+ T Cell Isolation Kit (Stemcell Technologies, #17953) through negative selection. These isolated cells were then cultured in RPMI 1640 medium (ThermoFisher, #11875093) containing 10% fetal bovine serum (FBS) (ThermoFisher, #10437028), 100 U/mL penicillin, 100 μg/mL streptomycin (ThermoFisher, #15140122), and 100 U/ml human IL-2 (Miltenyi Biotec, #130-097-743) at 37°C in a humidified atmosphere of 5% CO_2_. To measure the activity of type-I interferon in the supernatant of infected and transfected CD8 T cells, a reporter monocytic human cell line called THP1 dual knockouts (TBK1^-/-^ IKKα^-/-^ IKKβ^-/-^ IKKε^-/-^) was used. This cell line lacks the expression of TBK1, IKKα, IKKβ, and IKKε, but not interferon signaling, and was cultured at 37°C in humidified atmosphere of 5% CO_2_ in RPMI 1640 medium supplemented with 10% of fetal bovine serum, 100 U/mL penicillin, and 100 μg/mL streptomycin.

### Proliferation of CD8 T cells

U-shaped 96-well plates were coated overnight at 4°C with 100 µl of 2mg/ml anti-CD3 (BD Bioscience, #555336) and 2mg/ml anti-CD28 antibodies (BD Bioscience, #555725) in PBS. As a control, cells were cultured in wells that contained only PBS overnight. On the following day, purified CD8 T cells were stained with CellTrace™ Violet Cell Proliferation dye (1µg/ml) for 15 min, then washed and resuspended in fresh RPMI media containing 10% FBS, 100 U/mL penicillin, 100 μg/mL streptomycin, 100 U/mL human IL-2, and 20 ng/mL IL-15. After that, the antibody cocktails were removed from the coated wells, and 1×10^5^ purified CD8 T cells were added to each well. The cells were then cultured for 3 days at 37°C in a humidified atmosphere of 5% CO_2_. When cell counting was needed, 20,000 beads were added before flow cytometry acquisition.

### IAV infection and RIG-I stimulation of CD8 T cells

CD8 T cells were exposed to the reassortant Influenza A virus (IAV) strain from PR8 and A/Brazil/11/1978: RG-PR8-Brazil78 HA, NA (H1N1) at a multiplicity of infection of 10 in serum-free medium. After one hour, the cells were washed twice with PBS and then incubated for varying durations depending on the experiment. RIG-I ligands were generated using a Transcript Aid T7 in vitro transcription kit (ThermoFisher, #K0441) with annealed DNA oligonucleotides serving as a dsDNA templates, as previously described (50) (sequence: TTGTAATACGACTCACTATAGGGACGCTGACCCAGAAGATCTACTAGAAATAGTAGATCTTCTGGGTCAGCGTCCC). A single stranded 3p-RNA, was generated using the dsDNA template (sequence: CGCGCGTAATACGACTCACTATAGGGAGCGCAGACGCGAGCGCGGCACGGCCGCCAAGGCGAGAC) and used as a negative control for RIG-I receptor activation. Lipofectamine 2000 (Thermo Fisher Scientific, Langerwehe, Germany, #11668019) was used to transfect RIG-I ligands. 5 µg/mL RNA and 2.5 µL/mL lipofectamine 2000 were mixed together and allowed for 15 mins to form a complex. After that, the mixture was added to 10^6^ NK cells/mL for the stimulation. In some experiments, recombinant IFN-α2a (1000 U/mL) was used to stimulate the cells (Miltenyi Biotec, Bergisch Gladbach, Germany, #130-108-984).

### Western blot

After harvesting, cells in equal counts were centrifuged at 500xg for 5 minutes and washed with PBS. The cells were then lysed with 1x Laemmle buffer, which included PhosStop (Roche, #4906837001) and protease inhibitor (Roche, #4693116001). The lysate was vortexed and incubated at 95°C for 5-7 minutes in a thermomixer, shaking at 600 rpm, to denature the bound proteins. The lysate was loaded into a 10% SDS-PAGE gel for electrophoresis until desired separation was achieved. The proteins were then transferred to nitrocellulose membranes and stained individually using a sequential staining approach with primary antibodies. The primary antibodies used were anti-β-actin mouse mAb (LI-COR Bioscience, #926-42212), anti-IFIT1 rabbit mAb (Cell Signalling, #14769), anti-phospho-p65 rabbit mAb (Cell Signalling, #3033), anti-phospho-TBK1 rabbit mAb (Cell Signalling, #5483), anti-RIG-I rabbit mAb (Cell Signalling, #3743), anti-STAT2 rabbit mAb (Cell Signalling, #72604) & anti-phospho-STAT2 rabbit mAb (Cell Signalling, #88410). After staining, the proteins were imaged using the Odyssey Imaging system (LI-COR Biosciences). The Image Studio Lite software was used to quantify the relative expression levels of the target protein by normalizing the signal intensity of the target protein to the signal intensity of β-actin.

### Flow cytometry and degranulation assay

Cells were rinsed with FACS buffer consisting of 2% FBS, 0.5 µM EDTA in PBS, and then incubated at 4 °C with anti-hCD3 BV-510 (BD Bioscience, #563109), anti-hCD8 APC (Miltenyi Biotec, #130-110-679) or FITC (BD Bioscience, #561948), anti-hCD69-BV650 (BD Bioscience, #563835), anti-hCD107a-PE (BD Bioscience, #555801) antibodies, anti-hCD25-PE (BD Bioscience, #555432), and Fixable Viability Dye eFluor™ 780 (Thermofisher, #65-0865-14) for 30 minutes. To perform the degranulation assay, 50,000-100,000 CD8 T cells were stimulated with anti-hCD3 (2µg/ml) and anti-hCD28 (2µg/ml) coated for 4 hours at 37°C. During the entire assay time, Golgi Stop (BD Bioscience, #554724), Golgi Plug (BD Bioscience, #555029), and anti-hCD107a-PE antibodies were added to the media. For intracellular protein staining, cells were fixed, permeabilized with eBioscience™ Foxp3 / Transcription Factor Staining Buffer Set (Thermofisher, #00-5523-00), and then incubated with anti-hIFN-γ-AF700 (BD Bioscience, #557995), anti-hTNF-PECy7 (BD Bioscience, #560678), and NP-FITC (Abcam, #ab210526) for 30 minutes at 4 °C. Gating was performed for CD8 T cell based on the expression of CD3 and CD8. CD8 cells were defined as CD3+ CD8+ cells, and the purity was determined to be greater than 90% for every donor (Suppl. Fig. 1).

### IFN-I reporter assay

The type-I IFN reporter activity was determined by collecting cell-free supernatants 16-20 hrs after infection or stimulation. 100 µl of the supernatant was then added to medium-free THP1-dual TBK1^-/-^ IKKα^-/-^ IKKβ^-/-^ IKKε^-/-^ cells and incubated for 24 hrs. To measure luciferase activity, 30 µl of the supernatant was mixed with 30 µl of a water solution of coelenterazine (1 µg/ml) in a white 96-well F-bottom plate, and the activity was measured immediately using an EnVision 2104 Multilabel Reader device.

### CRISPR-editing of primary CD8 T cells

We used pre-designed crRNAs to modify genes in primary human CD8 T cells utilizing the CRISPR/Cas9 system. To target the RIG-I gene (also known as DDX58), we used two crRNAs that aimed at the negative strand (GGATTATATCCGGAAGACCC) and the positive strand (GATCAGAAATGATATCGGTT). Moreover, we employed a pre-designed crRNA that directed the Cas9 endonuclease enzyme to cut the positive strand of STAT2 (AAGTACTGTCGAATGTCCAC) specifically. Electroporation was carried out using the P3 Primary Cell 4D-Nucleofector X Kit S (Lonza, # V4XP-3032) and the 4D Nucleofector system (Lonza) with the EH-115 program. We used up to 1.5X10^6^ primary human CD8 T cells per reaction to ensure the maximum viability and uptake of the CRISPR/Cas9 mixture. The CRISPR/Cas9 mixture was prepared by combining 2 µL of 100 µM crRNA for each crRNA, equal amounts of tracrRNA, 1.7 µL of Cas9 enzyme, and 1 µL of 100 µM enhancer. The mixture was completed to 25 µL per reaction with electroporation buffer (ratio of 16.65 µL P3 media to 3.65 µL supplement). After electroporation, the cells were incubated for 3 days before stimulation and infection.

### Statistical analysis

The statistical calculations were executed using GraphPad Prism 9, and each donor is distinguished by a single dot in all figures, with various colors to identify between donors. The bars signify the mean ± standard error of the mean (SEM) across all donors. Paired t-tests were employed to determine the significance of differences between two groups, while repeated measures one-way ANOVA followed by Dunnett’s correction was used for more than two groups. For multiple comparisons, two-way ANOVA followed by Bonferroni’s correction was used. Statistical significance was denoted by asterisks as follows: *p < 0.05, **p < 0.01, ***p < 0.001, and ****p < 0.0001.

## Author Contributions

A.A.M. performed the majority of the experiments, while C.W. and C.H. contributed by performing some of the experiments. A.A.M., G.H., S.S., A.G.B. and M.S. conceived experiments. A.A.M., S.S., A.G.B. and M.S. analyzed and interpreted data and wrote the manuscript. Funding acquisition: G.H., A.G.B. and M.S. All authors revised the manuscript. All authors have read and agreed to the published version of the manuscript.

## Institutional Review Board Statement

Not applicable.

## Informed Consent Statement

Buffy coats were collected from anonymous healthy donors with written informed consent, in compliance with the principles of the Declaration of Helsinki, and approved by the Ethics Committee of the Medical Faculty, University of Bonn (approval no. 516/20).

## Data Availability Statement

Not applicable.

## Acknowledgments

We would like to acknowledge the assistance of the Flow Cytometry Core Facility at the Institute of Experimental Immunology, Medical Faculty at the University of Bonn—Project number 216372545.

---

**Supp. Fig. 1.**
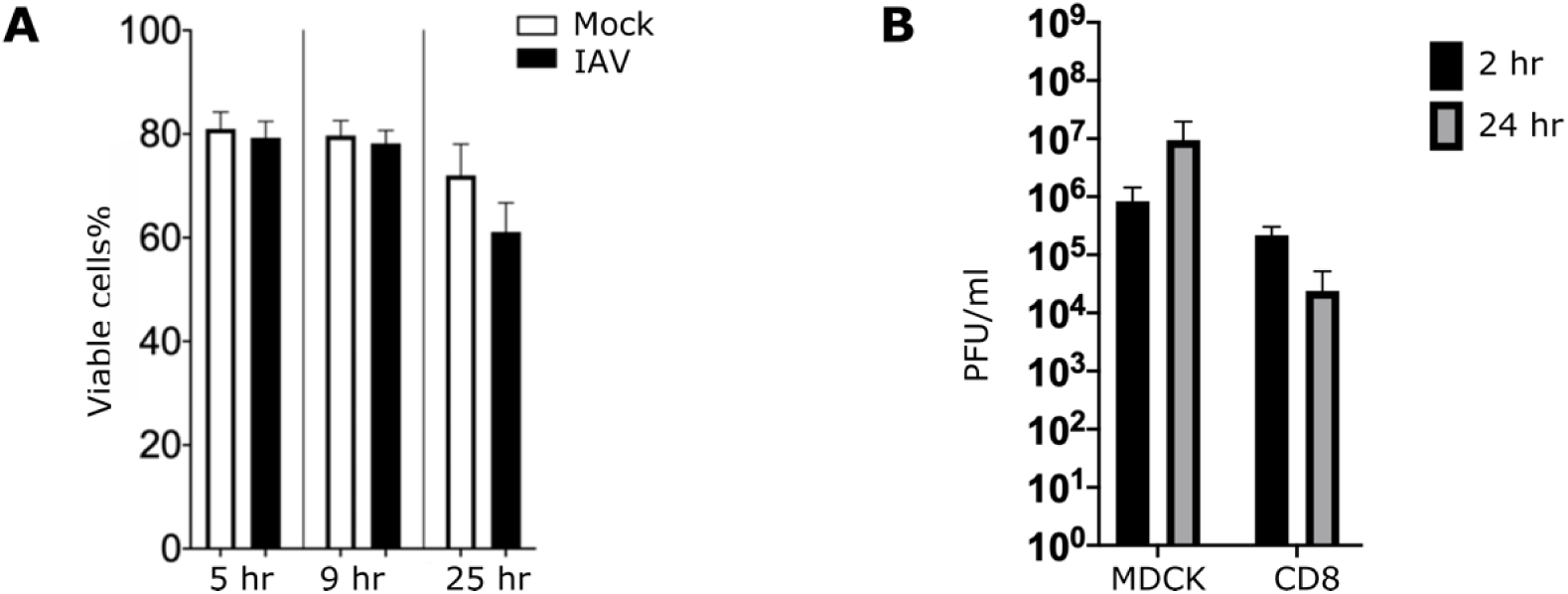
Infection outcomes of IAV in CD8 T cell. Two bar charts showing the results of experiments investigating the effects of influenza A virus (IAV) infection on CD8 T cells. **(A)** a bar chart representing the viability of CD8 T cells mock-infected or infected with IAV at different time points (5, 9, and 25 hrs post-infection). **(B)** display the viral titer (PFU/ml) measured in cell-free supernatants collected from CD8 T cells and MDCK cells positive control at 2 and 24 hrs post-infection (n= 5, mean ± SEM).

**Supp. Fig. 2.**
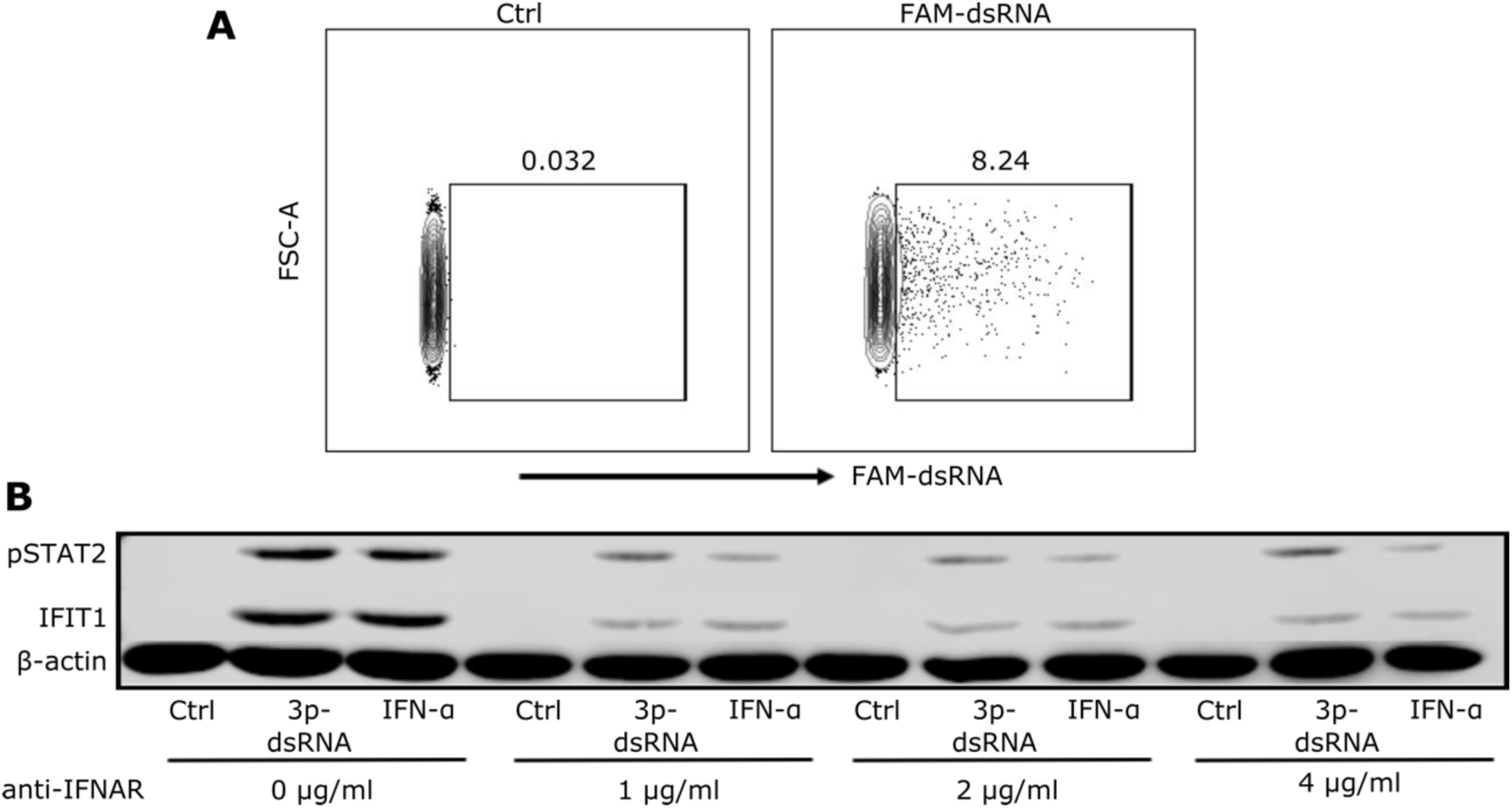
3p-dsRNA induces type-I IFN secretion and stimulates IFIT1 in an IFNAR dependent mechanism. **(A)** Flowcytometry plot showing CD8 T cells treated with unlabeled 3p-ssRNA or FAM-labeled 3p-dsRNA (FAM-dsRNA) **(B)** Western blot image for CD8 T cells either pre-treated without anti-IFNARα antibodies or with 1, 2, or 4 µg/ml for one hr then control RNA (Ctrl), 3pdsRNA, or IFN-α were added. Phospho STAT2 and IFIT1 proteins were used to detect the activation of IFNAR.

**Supp. Fig. 3.**
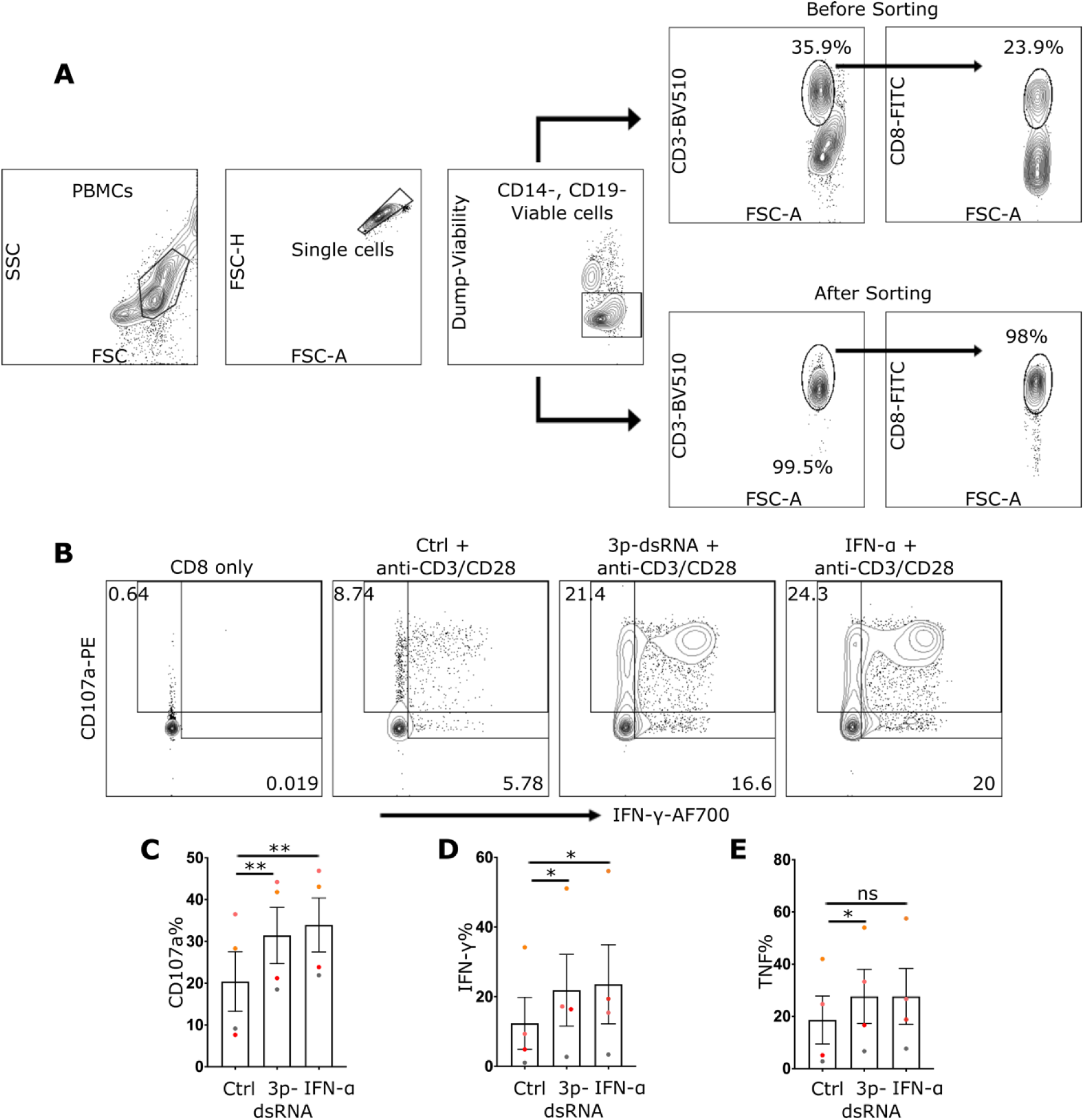
Sorted CD8 T cells reproduced the results from using purified CD8 T cells. **(A)** Representative flowcytometry plots showing sorting strategy and the purity of CD8 T cells before and after sorting. **(B)** Illustrative flowcytometry plot for sorted CD8 T cells treated with media only, or Ctrl RNA, 3p-dsRNA or IFN-α then stimulated with anti-CD3/CD28 antibodies. The plots demonstrate CD107a against IFN-γ or TNF against CD69 respectively. **(C-E)** quantification of the previously mentioned activation markers. Every donor is represented by a colored dot, bars show mean ± SEM. Repeated measures one-way ANOVA followed by Dunnett’s correction for more than two groups (ns= not significant, *p < 0.05, and **p < 0.01).

**Supp. Fig. 4.**
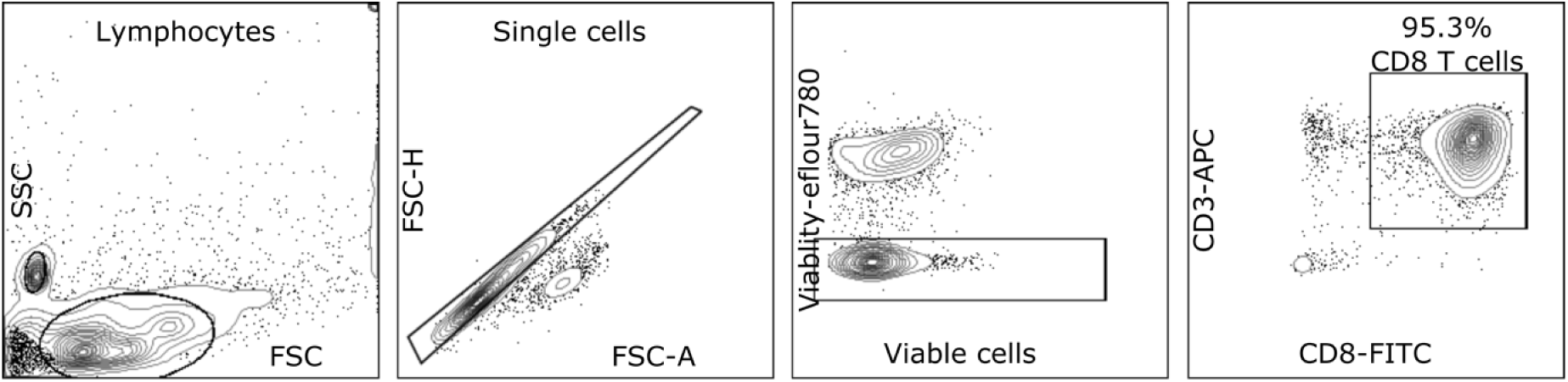
Gating strategy and CD8 T purity. Representative flow cytometry plots of peripheral blood mononuclear cells (PBMCs) obtained from one of the donors. The lymphocyte population was initially gated based on their side and forward scatter characteristics. We then used negative gating to exclude dead cells, and CD8 T cells were identified as cells expressing both CD3 and CD8 markers.

